# Into-India or out-of-India? Historical biogeography of freshwater gastropod genus *Pila* (Ampullariidae)

**DOI:** 10.1101/643882

**Authors:** Maitreya Sil, N. A. Aravind, K. Praveen Karanth

## Abstract

The biota of the Indian subcontinent has assembled during various points of the history of its continental drift: some when it was still a part of Gondwanaland and subsequently dispersed ‘out-of-India’ and some dispersed ‘into-India’ after it collided with Asia. However, the relative contribution of these connection to the current biotic assembly of the subcontinent is still under-explored. We aimed to understand the relative importance of these various routes of biotic assembly in India through studying the historical biogeography of tropical Old World freshwater snail genus *Pila*. We reconstructed a near-complete phylogeny of Ampullariidae including all the described Pila species from India and published sequences of Ampullariids from all over the world from two mitochondrial and two nuclear markers. Thereafter molecular dating and ancestral area reconstruction analyses were carried out in order to ascertain the time frame and route of colonization of India. The results suggest that Pila dispersed into India as well as other parts of tropical Asia from Africa after both India and Africa collided with Eurasia. Furthermore, multiple dispersals have taken place between Southeast Asia and India. The findings consolidate the rapidly building evidence that much of the current assemblage of biota actually dispersed into-India after it collided with Asia.

## 1. Introduction

The current assembly of flora and fauna is an outcome of a combination of present and historical processes (Brown & Lomolino, 1998). Geological processes such as continental drift is a major historical process driving floral and faunal assemblages. In this regard, the Indian subcontinent is of much interest to historical biogeographers owing to its plate tectonic history. India was a part of Gondwanaland supercontinent along with Africa, South America, Antarctica, Australia and Madagascar. Indo-Madagascar was separated from the rest around 121 million years ago (mya) (Sanmartín et al., 2004). Later, India broke off from Madagascar around 80 mya (Chatterjee and Scotese, 1999), drifted northwards and came in contact with Asia around 55–35 mya (Aitchison, Ali, & Davis, 2008). The complex geological history of India thus gives us an opportunity to understand how plate tectonic processes shape assembly of biota.

Mani (1974) classified Indian biota into several categories based on their purported origin such as Ethiopian (African) elements in the west, Palearctic elements in the north, Indo-Chinese or Sundaic (Southeast Asian) elements in the east and endemic Peninsular elements in the South. However, the origins of the Peninsular forms are much debated. It was speculated and later established that some of them were present in peninsular India (PI) and Sri Lanka since the Indian plate broke off from Madagascar around 90 million years ago (mya) such as the endemic frog family Nasikabatrachidae (Biju & Bossuyt, 2003) and while other lineages dispersed into SEA ‘out-of-India’ as well: Ichthyophiid caecilians (Gower *et al.*, 2002), Dipterocarpaceae (Dayanandan *et al.*, 1999) and Crypteroniaceae (Conti *et al.*, 2002). Nevertheless, some of the forms found in wet evergreen forests of the Western Ghats in PI were hypothesized to have sister/closely related forms in Northeast India and Southeast Asia (Mani 1974). Indeed, molecular phylogenetic studies have showed that many peninsular Indian forms, not exclusively wet zone taxa, indeed have their origins in Southeast Asia (Honda *et al.*, 1999; Noonan & Chippindale, 2006; Köhler & Glaubrecht, 2007; Van Bocxlaer *et al.*, 2009; Adamson, Hurwood, & Mather, 2010; Jansen, Savolainen, & Vepsäläinen, 2010; Rehan *et al.*, 2010; Datta-Roy *et al.*, 2012; Agarwal & Karanth, 2015; Barley *et al.*, 2015; Klaus *et al.*, 2016). There are certain theories that speculate that during India’s northward course some groups dispersed into the Indian plate from Africa, when it lay close to the latter (Briggs, 2003). These dispersals have also been attributed to a land bridge, in the form of a chain of islands called the Oman-Kohistan-Dras Island Arc, connecting the two landmasses around 75–65 mya (Chatterjee and Scotese, 2010). Furthermore, once the African plate sutured with Eurasia around 40 mya (Van Yperen, 2005), a host of diverse African elements dispersed into India through the ‘Eurasian route’. Thus, Indian biota has diverse origins, some have ancient Gondwanaland affinity while others have dispersed more recently from Asia or Africa ‘into-India’ (Praveen Karanth, 2006; Datta-Roy & Praveen Karanth, 2009). However, the relative importance of these different routes of faunal assembly in India is still not well understood. Further investigation into understudied groups found in Peninsular India which have African as well as SE Asian affinities will provide better insight into the patterns of faunal assemblages over time in Peninsular India.

The signatures of continental drift on biogeographic events are best observed in freshwater systems largely owing to their mode of dispersal: along the flow of water. Physical link among the freshwater bodies between different landmasses requires terrestrial connection. This ensures most of the dispersals undertaken by freshwater organisms would be geodispersal events. Hence, their phylogenies often mirror continental break ups and connections. The genus *Pila* is distributed in tropical and sub-tropical Asia including India, Africa and Madagascar. It belongs to the family Ampulariidae which is found in many of the Gondwanan landmasses such as: India, Africa, Madagascar and South America as well as in Central America and Southeast Asia (SEA). The presence of *Pila* in SEA, Madagascar and Africa makes it an interesting model system to ascertain the role drifting Indian plate on the assembly of SEA biota. Previously it was speculated that the Indian members of *Pila* are an Gondwanan element that was present in PI since the fragmentation of the supercontinent and was carried to Asia by drifting Indian plate (Berthold, 1991) acting as a ‘biotic ferry’. The biotic ferry hypothesis has been put forward and supported previously by various a number of studies on various taxonomic groups (Mani, 1974; Hedges, 2003; Bossuyt et al., 2006). However, this scenario is not supported by limited molecular phylogenetic study undertaken on this group (Hayes, Cowie, & Thiengo, 2009). In that study the split between African and Asian *Pila* is much more recent than the split between old world and new world Ampullariids, which is contrary to the order of fragmentation of Gondwanaland. Hence, they concluded that *Pila* colonized Asia later after Africa collided with Eurasia. Nevertheless, we believe this result is tentative as the sampling from Asia was sparse: especially no species from India were included and molecular dating analysis was not carried out. Given that India has ancient Gondwanan links, incorporating species from India might alter the outcomes of the story. Assuming that Gondwana origin hypothesis is rejected then there are additional scenarios that need to be tested : 1) *Pila* colonized India from Africa when there was a brief connection between the two drifting landmasses (∼75–65 mya) and 2) The ‘Eurasian route’ hypothesis which postulates dispersal of African fauna into tropical Asia after the former collided with Eurasia (∼40 mya).

To this end, we carried out thorough sampling of *Pila* throughout the country to capture all the described species. Molecular phylogenetic, molecular dating and ancestral range estimation analysis were undertaken to ask the following questions: 1) Where did the genus *Pila* originate? 2) Did Indian lineages come ‘into India’ from SEA or Africa after India collided with Eurasia or is it a lineage that existed in India ever since it was a part of Gondwanaland and then dispersed ‘out of India’? 3) When did *Pila* colonize India and rest of tropical Asia?

## 2. Materials and methods

### 2.1 Sample collection

Whole animal samples were collected from throughout India covering all the major river basins. The samples were preserved in absolute ethanol. Five species are described from within the political boundary of India: *Pila globosa, Pila virens, Pila saxea, Pila cf. nevilliana* and *Pila olea* (Rao, 1989). Out of these, the first three are widespread and the rest are data deficient and known from only their type locality (Rao, 1989). Since many widely distributed species might be species complexes with many cryptic species (Lajmi, Giri, & Karanth, 2016; Deepak & Karanth, 2017; Karanth, 2017), the three widespread species were collected from multiple locations of their distribution range. We also collected individuals from the type locality of data deficient species as a proxy for those species. Furthermore, a hill stream dwelling species of *Pila* was collected from the Northeast (NE) Indian state of Mizoram, bordering Myanmar. This species is referred to as *Pila* sp. 1 Throughout the manuscript. See Appendix A Table A1 for the complete list of species with their sampling locations and Appendix A Figure A1 for a map of sampling locations. Additionally, five *Pila* species from SEA and Africa, three African and two South American Ampullariid genera were included in the dataset whose sequences were obtained from the GenBank (see Appendix A Table A2).

### 2.2 Molecular work

DNA was extracted using the CTAB extraction method from foot muscle tissue of individuals (Williams, Reid, & Littlewood, 2003, Chakraborty *et al*., under review). Two mitochondrial (COI and 16S rRNA) and two nuclear markers (18S rRNA and Histone H3) were amplified. Sequencing was carried out at Medauxin Inc, Bangalore, India. Sequences were aligned thereafter using ClustalW in MEGA 7 (Kumar, Stecher, & Tamura, 2016). See Table 1 for a complete list of molecular markers used.

**Table 1.** Details of molecular markers used in the study.

| Gene name | Abbreviation | Primers | References |
| --- | --- | --- | --- |
| Cytochrome c oxidase subunit 1 | COI | LCO and HCO | (Folmer *et al.*, 1994; Williams *et al.*, 2003) |
| 16S ribosomal RNA | 16S rRNA | 16Sar-L and 16Sbr-H | (Palumbi, S.R., Martin, A., Romano, S., Mcmillan, W.O., Stice, L., Grabowski, 1991) |
| 18S ribosomal RNA | 18S rRNA | 18SYLMFOR<br>18SYLMREV | (Stothard *et al.*, 2000) |
| Histone H3 | Histone H3 | H3F and H3R | (Colgan *et al.*, 1998) |

#### Phylogenetic analysis

The mitochondrial and nuclear dataset was analysed separately first to detect mito-nuclear discordance if any and then were concatenated and analysed together. The models of sequence evolution and the partitioning scheme was derived from partition analysis using BIC employed in PartirionFinder (Lanfear *et al.*, 2017) (see Table 2 for complete list of partitions). Maximum likelihood analysis was carried out in RAxML HPC 1.8.2 (Stamatakis, 2014) implemented in raxmlGUI 1.5 (Silvestro & Michalak, 2011). Ten independent chains were run along with 1000 thorough bootstrap replicates. MrBayes 3.2 (Ronquist & Huelsenbeck, 2003) was employed to carry out the Bayesian analysis. Two independent runs were undertaken for 1000000 generations and sampled every 500 generations. Lowering of the standard deviation of split frequency below 0.01 was taken as a proxy for convergence. Additionally, Tracer 1.5 (Rambaut, 2009) was employed to see whether all the parameters have reached a stationary phase (>200 ess values). The phylogeny reconstructed from the combined dataset resembled the mitochondrial dataset more than the nuclear dataset. It is preferable to put more trust on the nuclear topology especially when one is investigating deeper divergences because mitochondrial genes evolve faster than nuclear genes and thus reach saturation sooner than the latter (Brown, George, & Wilson, 1979; Lajmi *et al.*, 2018). Especially, the of the third codon position of the mitochondrial coding genes reach saturation faster and are responsible for the discordance between mitochondrial and nuclear topology (Lajmi *et al.*, 2018). Hence, we excluded the third position from all the codons of the COI gene. The concatenated dataset thus prepared resembled the nuclear topology. Hence, the phylogenetic reconstruction, divergence dating analyses were carried out based on the concatenated dataset excluding the third codon position of COI.

**Table 2.** Details of the partition scheme used for each analysis.

|  | RAxML | MrBayes | BEAST |
| --- | --- | --- | --- |
| COI | GTR+I+G | GTR+I+G | TrN+I+G |
| 16S rRNA | GTR+I+G | HKY+I+G | TVM+I+G |
| 18S Rrna+ Histone H3 | GTR+I+G | K80+G | K80+G |

### 2.3 Topology tests

An Approximately Unbiased (AU) test was undertaken to establish whether the topologies retrieved from the ML analysis were significantly better than other topologies which could potentially support other biogeographic scenarios. AU test perform better than other such topology tests in controlling type 1 error (Shimodaira, 2002). Thus we carried out two additional ML analysis in raxmlGUI 1.5 using the same settings as the previous one, where the African and the other where the SE Asian species of *Pila* were constrained to be sisters the other *Pila* species respectively (see Appendix A Figure A2). Persite likelihood of all the three trees were estimated in PAUP 4b10 (Swofford, 2001). P-values for the AU test were calculated in CONSEL (Shimodaira & Hasegawa, 2001) utilizing the per-site likelihood values.

### 2.4 Molecular dating

Molecular dating was carried out using BEAST 1.8.3 (Drummond *et al.*, 2012) with the concatenated dataset. The models of sequence evolution and the partitioning scheme were followed as per the results from partition analysis. Uncorrelated lognormal relaxed clock was used for each partition with CTMC rate reference prior. A yule tree prior was used as the tree prior. The tree was calibrated using two old world Ampullariid fossils. A) The most recent common ancestor (MRCA) of the genus *Pila* was calibrated with the oldest fossil of *Pila* unearthed from Oman and dates back to Priabonian stage (37.8–33.9 mya) of Eocene epoch. B) Similarly, the MRCA of the genus *Lanistes* was calibrated with the oldest *Lanistes* fossil described from Maastrichtian (Late Cretaceous, 72.1–66 mya) deposits of Oman. The fossil calibration can only inform us about the minimum bound of the calibration with certainty. Hence, we set gamma priors on both the calibrations. Different combinations of the shape (α) parameter were employed to incorporate the uncertainty about the mode of the distribution in four independent runs, whereas the scale (β) paarmeters were fixed at 2 for both the calibrations (see Appendix B for more details). Thereafter, marginal likelihood score of the analyses were compared using Bayes factor test in Tracer v1.5. One final analysis was run for 100 million generations with sampling every 1000 generations using the best calibration scheme picked (*Pila* MRCA α parameter value=4, *Lanistes* MRCA α parameter value=4; *Pila* MRCA β parameter value=2, *Lanistes* MRCA β parameter value=2). We carried out unconstrained as well as constrained analyses with the same parameter settings in order to generate a chronogram to test an alternate biogeographic scenario. The AU test suggested that the tree in which the African species of *Pila* were constrained to be sisters to all the Asian *Pila* species was not significantly different and thus could not be rejected. Hence, in the second dating analysis the African species of *Pila* were constrained to be sisters to all the Asian *Pila* species. The saturation (ess>200) was checked in Tracer v1.5 (Rambaut, 2009). The tree was summarized in TreeAnnotator 1.8.0 (Rambaut, & Drummond, 2013).

### 2.5 Ancestral range estimation analysis

Ancestral range estimation analyses were carried out in R 3.4.2 (available at https://www.r-project.org/) using the package BioGeoBears (Matzke, 2013). DEC+*j* model was employed in order to find out the range evolution and time of colonization of India. DEC+*j* model has often been implemented to explore the range evolution of island taxa and dispersal limited taxa. (Kitson *et al.*, 2018; Hendriks *et al.*, 2019). In this particular case, freshwater snails are known to be dispersal limited. Moreover, many of the landmasses where the genus *Pila* is distributed in did not have land connection with each other for much of their geological history. Although, geodispersal is more likely for freshwater species, we cannot a priori rule out possibility of jump dispersal. Hence, we decided to implement the DEC+*j* model as it fulfils all the requirements. The Maximum Clade Credibility (MCC) tree, obtained from BEAST analysis, was pruned using the package APE (Paradis, Claude, & Strimmer, 2004) in R 3.4.2 to include only old world taxa. The distribution range of the family were broken up into 1) Africa, 2) SEA including NEI, 3) India covering all of India south of the Himalayas except NEI which is part of Indo-Chinese subregion) (Wallace, 1876). We carried out time stratified analyses where the adjacency and dispersal probability between different landmasses changed over time based on the plate tectonic movement of the said landmasses. Dispersal from Africa to India and vice versa was assigned high probability (1.0) at certain points of time and low probability (0.01) at other times as their adjacency changed over time. Likewise, different values were assigned to the dispersal between SEA and India and their adjacency as the relative position of these landmasses evolved over time. Direct dispersal from SEA to Africa was unlikely for most of their geological history because these landmasses were almost never in close proximity to each other. Hence dispersal between these two landmasses was given low probability (0.01) for most of the time periods, following previous studies where similar dispersal probability was assigned between areas where direct dispersal was less likely (Barley *et al.*, 2015) (see Appendix A Table A3 for details on the dispersal multiplier and adjacency between areas). Two analyses were carried out using two different MCC trees: 1) One analysis was carried out based on the ML topology. 2) Similarly, another analysis was carried out based on one of the constrained tree topologies, which was not significantly different from the ML tree.

## 3 Results

### 3.1 Phylogenetic analysis

The Bayesian tree and ML tree (Figure 1), derived from the dataset excluding third codon position of COI gene, exhibited overall congruence. The old world and new world taxa formed two clades. African genus *Saulea* was sister to all other Old World taxa. *Pila* was monophyletic and sister to African genus *Lanistes*, this clade was in turn sister to African genus *Afropomus*. Within the *Pila* radiation, Peninsular Indian taxa *Pila saxea* was sister to all the remaining *Pila* species. The clade consisting of remaining *Pila* species constituted four lineages, these included 1) African species *P. speciosa* and *P. ovata*, 2) *P. gobosa* distributed in India and SEA, Peninsular Indian species *Pila virens, P.cf. nevilliana*, from Peninsular India and *P. olea*, from NEI, 3) *P. conica, P, polita, P. ampullaceal* from SEA, and *Pila* sp 1 from NEI. There were two well supported clades in the *P. globosa* group: one consisting of species sampled from Gangetic basin and the other was composed of individuals collected from Orissa and Northeast India. *P. nevilliana* was nested within *P. virens*. Hereafter this individual is referred to as *P. virens* 7. *P. olea* was retrieved as the sister to the *P. virens* clade. Although the bootstrap values strongly suggested sister relationship of *Pila* and *Lanistes* and monophyly of *Pila*, support for relationships within genus *Pila* was low.

**Figure 1.**
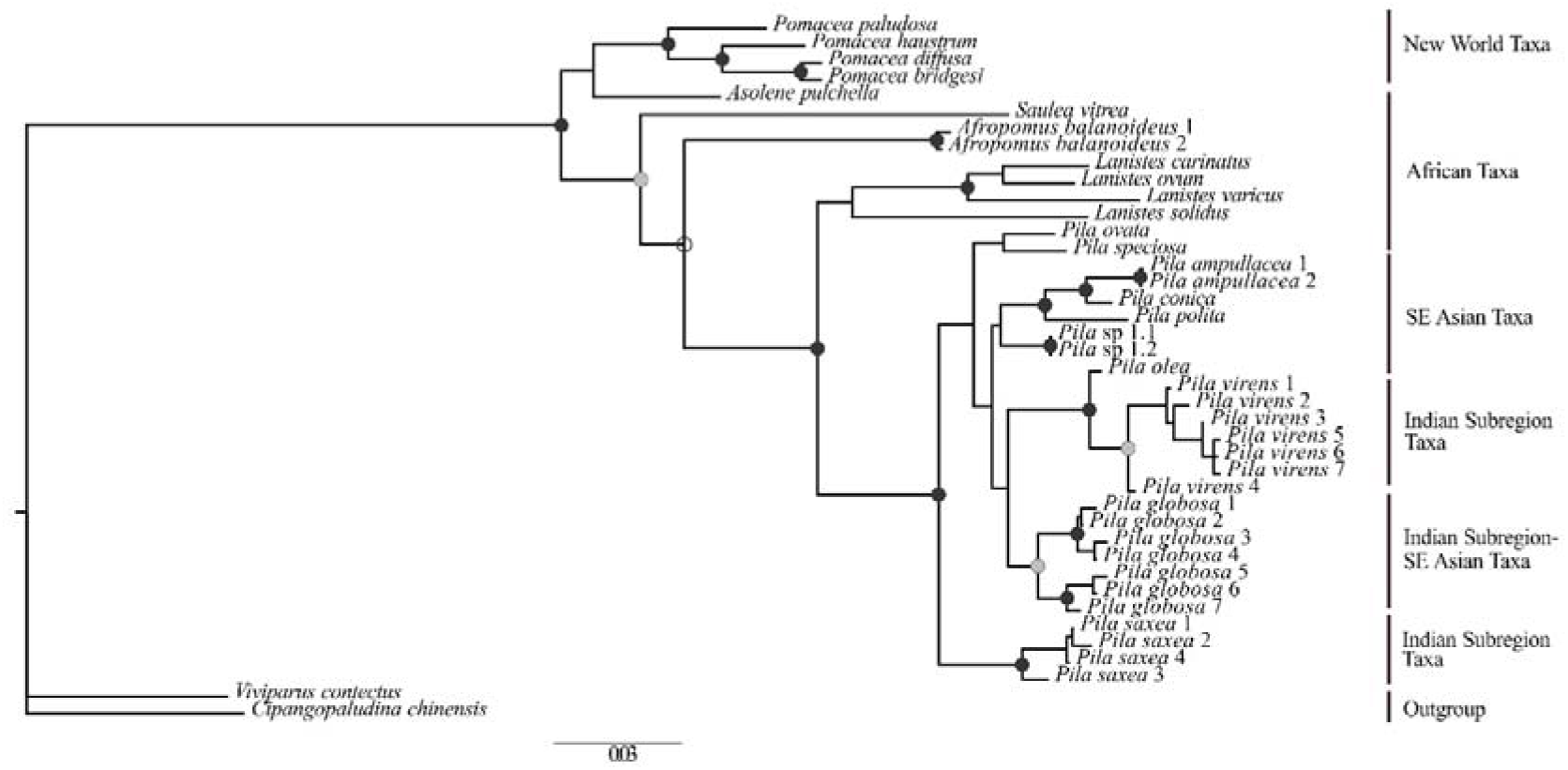
The Bayesian inference tree obtained from MrBayes analysis. The black circles represent nodes with high Bayesian posterior probability (>0.9) and high bootstrap support (>80), whereas, the green circles represent only high posterior probability.

### 3.2 Topology tests

The results of the AU test suggest that the topology where the African species of *Pila* were constrained to be sisters to all other *Pila* species was not significantly different from the best tree. The other topology however, where SE Asian species were constrained to be sister to the other *Pila* species, was significantly different and thus rejected, (P=0.008).

### 3.3 Molecular dating

The model where, the alpha parameter of the gamma distribution prior of both the MRCA of *Pila* and *Lanistes* were fixed as 4, was retrieved as the best fit model in Bayes factor test. Hence, we discuss results from only that particular analysis. The time tree (Figure 2.4) obtained was similar to the Bayesian and likelihood tree (figure 2). The split between old world and new world groups dated back to 188.2–107.1 mya. The split between *Pila* and *Lanistes* was found to be 99.6–69.2 mya. Radiation started in genus *Pila* at around 55–35.8 mya. The African species diverged from their sister lineage at around 43.3–25.8 mya. The SEA species, which were retrieved to be sisters to the clade containing *P. gobosa, P. virens*, and *P. olea*, split off from the latter at around the same time (38.7–22 mya). Both in African and SEA species radiation commenced immediately afterwards (33.3–3.7 mya and 35–17.5 mya respectively). The clade consisting of *P. globosa* branched off from the sister clade 34.1–17.6 mya. The two clades within the *P. globosa* group diverged around 22.1–7.6 mya. *P. olea* split off from the entire clade containing *P. virens* around 24.1–8.7 mya. The reconstruction in which the African species were sister to all other species of *Pila* (Appendix A Figure A3), divergence between African *Pila* and the rest dates back to 49.5–35.7 mya.

**Figure 2.**
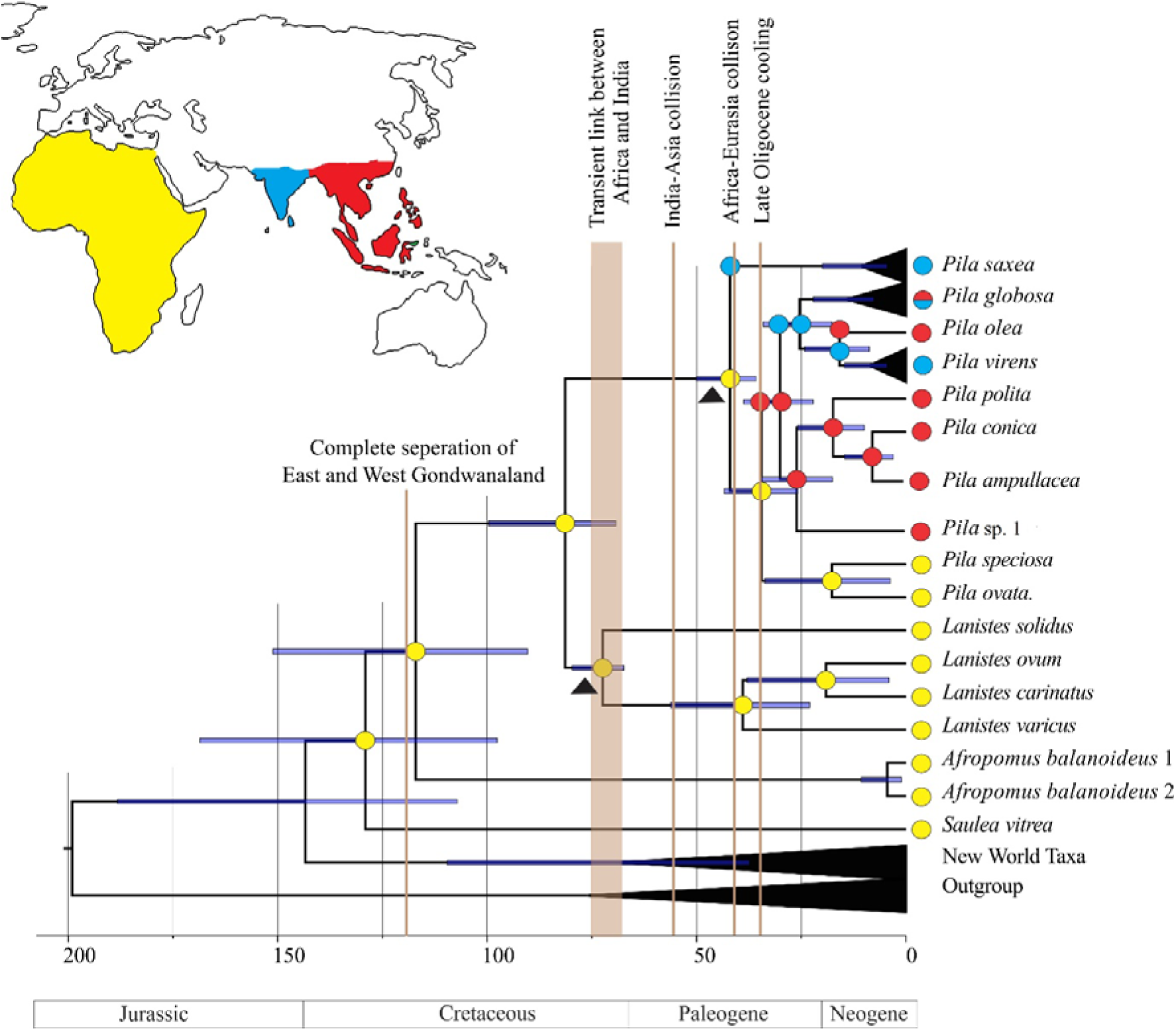
The MCC tree obtained from the BEAST analysis with estimated ancestral ranges mapped on the nodes. The yellow, blue and red circles stand for Africa, Indian subregion and SEA respectively, whereas the bicoloured ones indicate presence in any two of all the areas, depending on the constituent colours. The grey bars are the 95% highest posterior density estimates for the age of each node. The arrow indicate colonization of Indian subcontinent, while the black triangle show the placement of the geological prior

### 3.4 Ancestral range estimation analysis

The ancestral range estimation analyses using both the unconstrained and constrained MCC tree (Figure 2 and Appendix A Figure A3) obtained fairly similar results. The Old World Ampullariids as well as *Pila* originated in Africa as per expectation, given the higher genus level diversity in Africa. The Indian subcontinent was colonized from Africa after the Gondwanaland break up, which was speculated in previous studies (Hayes *et al.*, 2009). Furthermore, dispersals have taken place from SEA into India and back.

1. Results of the analysis using the unconstrained MCC tree (Figure 2) showed that the *P. saxea* lineage dispersed into India from Africa during 50–35 mya. The dispersal of the lineage leading to all other Asian *Pila* followed immediately afterwards (43–25 mya) from Africa into SEA. Soon after, the lineage consisting of *P. virens, P. globosa* and *P. olea* dispersed into India from SEA (38.7–22.1 mya). Two independent dispersals have taken place from India to SEA consisting of one jump dispersal and one range expansion events. *P. olea* dispersed into SEA during 24.1–8.7 mya followed by range expansion of *P. globosa* (34.1–0).
2. The results of the analysis using the constrained MCC tree (Appendix A Figure A3), unlike the pervious analysis showed that the lineage consisting of all Asian *Pila* not just *P. saxea* colonized India from Africa (45.9–35.7 mya). This was followed by a dispersal into SEA from India by the lineage composed of all extant Asian *Pila* species except *P. saxea* (45.2–27.3 mya). Soon after there was a back dispersal to India by a lineage consisting of *P. virens, P. globosa* and *P. olea* (39.2–22.3). Similar to the last analysis, *P. olea* dispersed into SEA during 24.1–8.7 mya followed by range expansion of *P. globosa* (35.2–0).

## 4 Discussion

The study aims to understand the range evolution of tropical Old World freshwater gastropod genus *Pila* especially the origin and biogeography of the Indian species. The results suggested that the genus has its origin in Africa. *Pila* colonized tropical Asia from Africa during Eocene-Oligocene times. Thereafter, multiple dispersals have taken place within tropical Asia: India and SEA. Below we discuss how the paleogeology and paleoclimate of the three colliding continents: Africa, Eurasia and India that could have governed these dispersal events.

### 4.1 From Africa into India?

Africa has shared a long biogeographic link with India. Even after Indo-Madagascar’s separation from Africa, there is evidence of a dispersal route between these continents. Geological and paleontological evidences suggest that India might have been connected to Africa by means a land bridge during late Cretaceous (Chatterjee and Scotese, 2010). However, a volcanic explosion at the end of Cretaceous, that led to the formation the Deccan traps, is thought to have wiped out much of the biota that was present in the subcontinent at that time (Officer *et al.*, 1987; Joshi & Karanth, 2011). Hence much of the biota that has African affinities probably colonized the subcontinent much later, after Africa collided with Eurasia.

The constrained and unconstrained tree topologies differed in their biogeographic inference. The analysis based on the unconstrained tree suggests that one dispersal from Africa to India leading to the *P. saxea* lineage took place during Eocene times (50–35 mya). The results further suggest that there was a second dispersal to SEA from Africa from mid-Eocene to mid-Oligocene (43–25 mya) which gave rise to all other extant Asian species. On the other hand, the analysis based on the constrained tree suggests that only one dispersal event occurred from Africa to India that gave rise to all extant Asian species during Eocene (50–35 mya). The analysis based on the constrained tree is more plausible given it suggests that only one dispersal took place from Africa into tropical Asia, yet, a more complete species and gene sampling has to be carried out to choose between the two scenarios. Although the number of dispersals is different between the two analyses, the directionality (from Africa into tropical Asia including India) and time frame of the dispersals are congruent. Incidentally this time frame coincides with collision of Afro-Arabian plate with Eurasian plate in late Eocene (∼40 mya) (Van Yperen, 2005).

The earliest fossil that resembles *Pila* was unearthed from Africa during late Cretaceous (Neubert et al., 2012). However, the assignment of this fossil to *Pila* genus is dubious. The earliest confirmed *Pila* fossil was excavated in Oman date back to late Eocene times. Thus, it is likely that the *Pila* lineages have followed a Eurasian route outlined by (Yoder & Yang, 2004). This is also concurrent with the closure of several major seaways connecting the Tethys with Atlantic and arctic oceans which would have dispersal across Eurasia possible (Ã & Sissingh, 2003). The results support the Eurasian route suggested for the dispersal of Loris from Africa although possibility of existence of such a route for dispersal of terrestrial biota at that period of time is contested by (Conti *et al.*, 2002). The time frame of the dispersal is also concurrent with completion of the suturing of the Asian and Indian plate (Aitchison *et al.*, 2008).

### 4.2 Role of Paleoclimate

*Pila* is primarily a lentic habitat species (Harzhauser *et al.*, 2016). Hence, humid climatic conditions that foster such habitats is likely to have aided in its dispersal. Oman in late Eocene was characterized by presence of vast swamplands which harboured Ampullariid snails such as *Pila* (Neubert & Damme, 2012). On the contrary present day Arabian Peninsula receive very less precipitation and host mainly arid habitats (Kotwicki & Al Sulaimani, 2009). The subsequent arid conditions in this region might be responsible for the absence of extant Ampullariid taxa in middle East. A major shift in global climate from being warm and humid to cold and arid took place around the Eocene-Oligocene boundary (∼34 mya) (Zachos *et al.*, 2001). The impact of these event on Central Asia and Arabian Peninsula is known from various sedimentological, biogeochemical, magnetostratigraphic and paleontological studies (Kraatz & Geisler, 2010; Sun & Windley, 2015). Our results from the unconstrained suggest that the dispersals from Africa into India and SEA took place during mid-Eocene to mid-Oligocene. The first dispersal by *P. saxea* lineage took place before the aforementioned drying event. The warm and humid climatic conditions present during that time period were likely to have aided in the first dispersal into India from Africa. It is plausible that the second dispersal was also facilitated by warm and humid climatic conditions in Eocene before the climatic shifts during Eocene-Oligocene boundary. Similarly, the only dispersal that took place from Africa into tropical Asia as suggested by the constrained analysis also occurred in Eocene before the commencement of the widespread aridification.

### 4.3 Dispersals across tropical Asia

Whilst, both the ancestral range estimation analyses favoured dispersal of *Pila* from Africa into tropical Asia (SEA and India) following the Eurasian route, the order of dispersal across different parts of tropical Asia were contrasting. Results from the ancestral range estimation analysis based on the unconstrained tree suggests a back dispersal of the lineage consisting of *P. virens, P. globosa* and *P. olea* into India from SEA (38.7–22.1 mya). The analysis implied two back dispersals to SEA as well. On the contrary, the other analysis showed that the lineage consisting of all the tropical Asia species excluding *P. saxea* dispersed into SEA, out of India (45.2–27.3 mya). This was followed by the dispersal of the lineage consisting of *P. virens, P. globosa* and *P. olea* into India (39.2–22.3 mya). *P. olea* dispersed back to SEA and *P. globosa* expanded its range into SEA soon after. A more complete sampling and inclusion of more molecular markers can help resolve this conflict. Interestingly, all these dispersal events took place after the completion of suturing of the Indian and Asian plates or overlapped with the event. It has been suggested before that there was an increase in dispersal rate from continental Asia to India after completion of the suturing (Klaus *et al.*, 2016). Hence, our study consolidates the finding that most of dispersals from continental Asia to India and vice versa occurred after the aforementioned event.

### 4.4 Rapid radiation

The genus *Pila* was monophyletic with high support, although, relationships within the clade were not well-supported. The reason behind the low bootstrap support can be attributed to the fact that different genes in the *Pila* lineage show different patterns of inheritance. The reason can be manifold: hybridization and random sorting of ancestral allele. Random sorting of ancestral alleles take place when more than one speciation events take place within a very short window of time and alleles are distributed randomly between these lineages such that the gene trees do not reflect the true species history (Whitfield & Lockhart, 2007; Rothfels *et al.*, 2012). Rapid radiation has been observed across the breadth of the taxonomic spectrum (Whitfield & Lockhart, 2007; Wiegmann, 2011; Rothfels *et al.*, 2012). In our study the very short internode distance hints towards a rapid radiation early in *Pila* lineage, which might have facilitated random sorting of ancestral allele, leading to the present issue. Rapid radiation can take place in response to increased ecological opportunity such as colonization of a new habitat with few species occupying similar niche.

### 4.5 Taxonomic implications

Individuals were collected from the type localities of the two data deficient species as a proxy for those species. Out of these two, *P. nevilliana* (*P. virens* 7) is nested within a *P. virens* radiations. It implies that perhaps the original collectors wrongly assigned it as a separate species. Indeed, some taxonomists expressed their reservations about whether this is a new species at all or a misidentified individual of *P. saxea* collected not from Tamil Nadu but Maharashtra (Annandale, 1921). Under any of these circumstances, the specimen included in the study is another *P. virens* individual. *P. olea* however was unrelated to all other *Pila* species collected from NEI. The original description also alludes to morphological similarities between these *P. olea* and *P. virens* (Prashad, 1924; Subba Rao, 1989). Incidentally, the lineage was sister to the clade containing *P. virens*. The molecular dating implies that the split dates back to 24.1–8.7 mya. Hence, it is likely that the individual does belong to *P. olea* and it does warrant a separate species status since it split off from its sister a long time back. However, it will require thorough morphological study before a conclusion is reached. The new species of Pila collected from NEI (*Pila* sp. 1) was initially thought to be *P. aperta*, a hill stream species of *Pila*, reported from Myanmar. However, careful observation of morphological characters suggest that this is actually not related to *P. aperta*, but possibly an undescribed species. Intriguingly, the two hill stream species: *P. saxea* and *Pila* sp. 1, are not monophyletic, suggesting independent evolution of hill stream dwelling traits. All the Indian species of *Pila* i.e. *P. globosa, P. virens* and *P. saxea* exhibited high genetic diversity. Exhaustive species delimitation analysis is needed to be carried out in order to ascertain whether these species are species complexes (Queiroz, 1998; Lajmi *et al.*, 2016; Deepak & Karanth, 2017; Karanth, 2017).

## 5. Conclusion

The study suggests an African origin of the genus *Pila*. According to the analysis based on the unconstrained tree: the ancestors of *P. saxea* dispersed to India from Africa, while the ancestors of *P. globosa* and *P. virens* together with the SE Asian species dispersed to SEA. The analysis based on the constrained tree suggests that a single dispersal event from Africa to India gave rise to all tropical Asian *Pila*. The time frame of dispersals, irrespective of the tree the analyses were based on, favours the Eurasian route theory which postulates dispersal of tropical Asian species from Africa after the latter docked with Eurasia. The ancestors of *P. globosa* and *P. virens* in turn colonized India from SEA. Gondwanan biota in India have colonized the subcontinent through different routes. Our study further confirms the finding that very few of these taxa were present in India when India was still part of Gondwanaland. Much of the biotic assembly in India occurred post-India-Asia collision through Eurasia.

## Acknowledgements

Anirudha Datta Roy, Ishan Agarwal and Aparna Lajmi commented on the manuscript, and assisted during analysis. Kunal Arekar, Chinta Sidhharthan, Harshal Bhosale, Pradip Barua, Tarun Singh, Shantanu Kundu, Suhel Sheikh, Animesh Mondal, Harshil Patel and Lansothung Lotha helped us during sample collection. The project was funded by DBT-IISc partnership program (22-0303-0007-05-469) to PK. The fieldwork was partially supported by Rufford Small Grant for Nature Conservation (19805-1) to MS.

## Appendix A

**Table A2.**
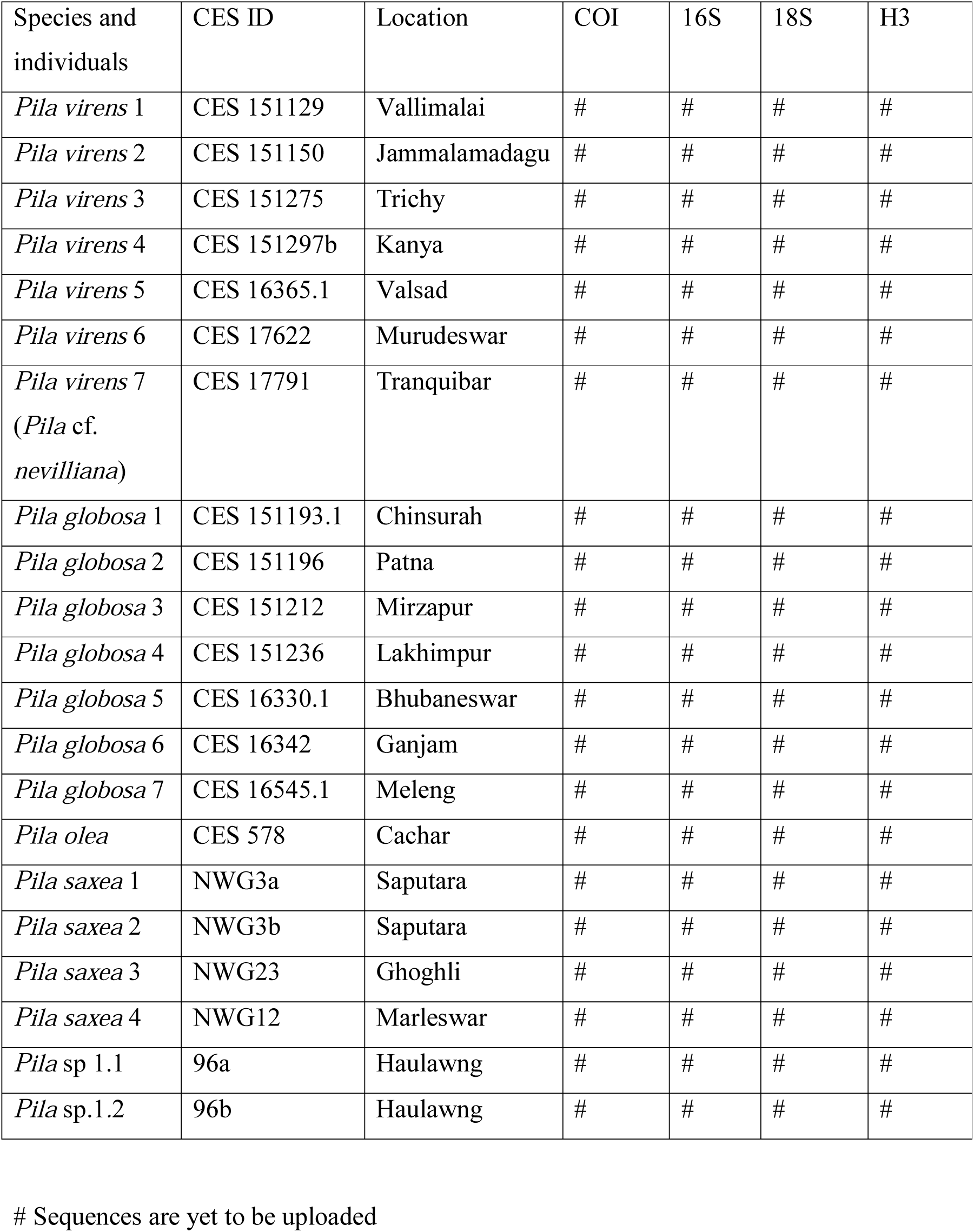
List of samples collected during the course of the study with their sampling location.

| Species and individuals | CES ID | Location | COI | 16S | 18S | H3 |
| --- | --- | --- | --- | --- | --- | --- |
| *Pila virens* 1 | CES 151129 | Vallimalai | # | # | # | # |
| *Pila virens* 2 | CES 151150 | Jammalamadagu | # | # | # | # |
| *Pila virens* 3 | CES 151275 | Trichy | # | # | # | # |
| *Pila virens* 4 | CES 151297b | Kanya | # | # | # | # |
| *Pila virens* 5 | CES 16365.1 | Valsad | # | # | # | # |
| *Pila virens* 6 | CES 17622 | Murudeswar | # | # | # | # |
| *Pila virens* 7<br>(*Pila* cf.<br>*nevilliana*) | CES 17791 | Tranquibar | # | # | # | # |
| *Pila globosa* 1 | CES 151193.1 | Chinsurah | # | # | # | # |
| *Pila globosa* 2 | CES 151196 | Patna | # | # | # | # |
| *Pila globosa* 3 | CES 151212 | Mirzapur | # | # | # | # |
| *Pila globosa* 4 | CES 151236 | Lakhimpur | # | # | # | # |
| *Pila globosa* 5 | CES 16330.1 | Bhubaneswar | # | # | # | # |
| *Pila globosa* 6 | CES 16342 | Ganjam | # | # | # | # |
| *Pila globosa* 7 | CES 16545.1 | Meleng | # | # | # | # |
| *Pila olea* | CES 578 | Cachar | # | # | # | # |
| *Pila saxea* 1 | NWG3a | Saputara | # | # | # | # |
| *Pila saxea* 2 | NWG3b | Saputara | # | # | # | # |
| *Pila saxea* 3 | NWG23 | Ghoghli | # | # | # | # |
| *Pila saxea* 4 | NWG12 | Marleswar | # | # | # | # |
| *Pila* sp 1.1 | 96a | Haulawng | # | # | # | # |
| *Pila* sp.1.2 | 96b | Haulawng | # | # | # | # |

**Table A2.** List of sequences downloaded from genbank from accession number.

| Species | Location | COI | 16S | 18S | H3 |
| --- | --- | --- | --- | --- | --- |
| **Outgroups** |  |  |  |  |  |
| *Cipangopaludina chinensis* | Haenam, S. Korea | EU528474.1 | FJ710213.1 | FJ710250.1 | - |
| *Viviparus contectus* | Denmark | EU274557.1 | EU274484.1 | EU274536.1 | EU274503.1 |
| **Species distributed in the New World** |  |  |  |  |  |
| *Pomacea paludosa* | Florida, USA | EU528476.1 | FJ710238.1 | FJ710279.1 | FJ710387.1 |
| *Pomacea huastrum* | Florida, USA | EU528479.1 | FJ710239.1 | FJ710280.1 | FJ710388.1 |
| *Pomacea diffusa* | Amazonas, Brazil | EU528480.1 | FJ710242.1 | FJ710283.1 | FJ710394.1 |
| *Pomacea bridgesi* | Laboratory Stock | EU528574.1 | EU274500.1 | EU274553.1 | EU274514.1 |
| *Asolene pulchella* | Buenos Aires, Argentina | EU528507.1 | FJ710225.1 | FJ710263.1 | FJ710371.1 |
| **Species distributed in Africa** |  |  |  |  |  |
| *Afropomus balanoidea* | Makemui, Sierra Leone | EU274558.1 | FJ710216.1 | FJ710253.1 | FJ710364.1 |
| *Lanistes carinatus* | Lake Bisinia, Uganda | EU274560.1 | EU274486.1 | EU274537.1 |  |
| *Lanistes ovum* | Lake Mweru, Zambia | EU274564.1 |  | EU274541.1 | EU274509.1 |
| *Lanistes solidus* | Chipanda, Malawi | EU274566.1 | EU274491.1 | EU274543.1 |  |
| *Lanistes varicus* | Mogtedo, Barkina Faso | EU274567.1 | - | EU274544.1 | EU274510.1 |
| *Pila ovata* | Uganda | EU274571.1 | EU274496.1 | EU274550.1 | - |
| *Pila speciosa* | Kinyasini, Zanzibar | EU274573.1 | - | EU274552.1 | - |
| **Species distributed in SEA** |  |  |  |  |  |
| *Pila ampullacea* | Binh Chu, Vietnam | EU528504.1 | FJ710218.1 | FJ710254.1 | - |
| *Pila polita* | Vietnam | EU528475.1 | FJ710220.1 | EU274551.1 | EU274513.1 |
| *Pila conica* | Hanoi, Vietnam | EU528588.1 | FJ710219.1 | EU274547.1 | EU274512.1 |

- No information could be found

**Table A3:**
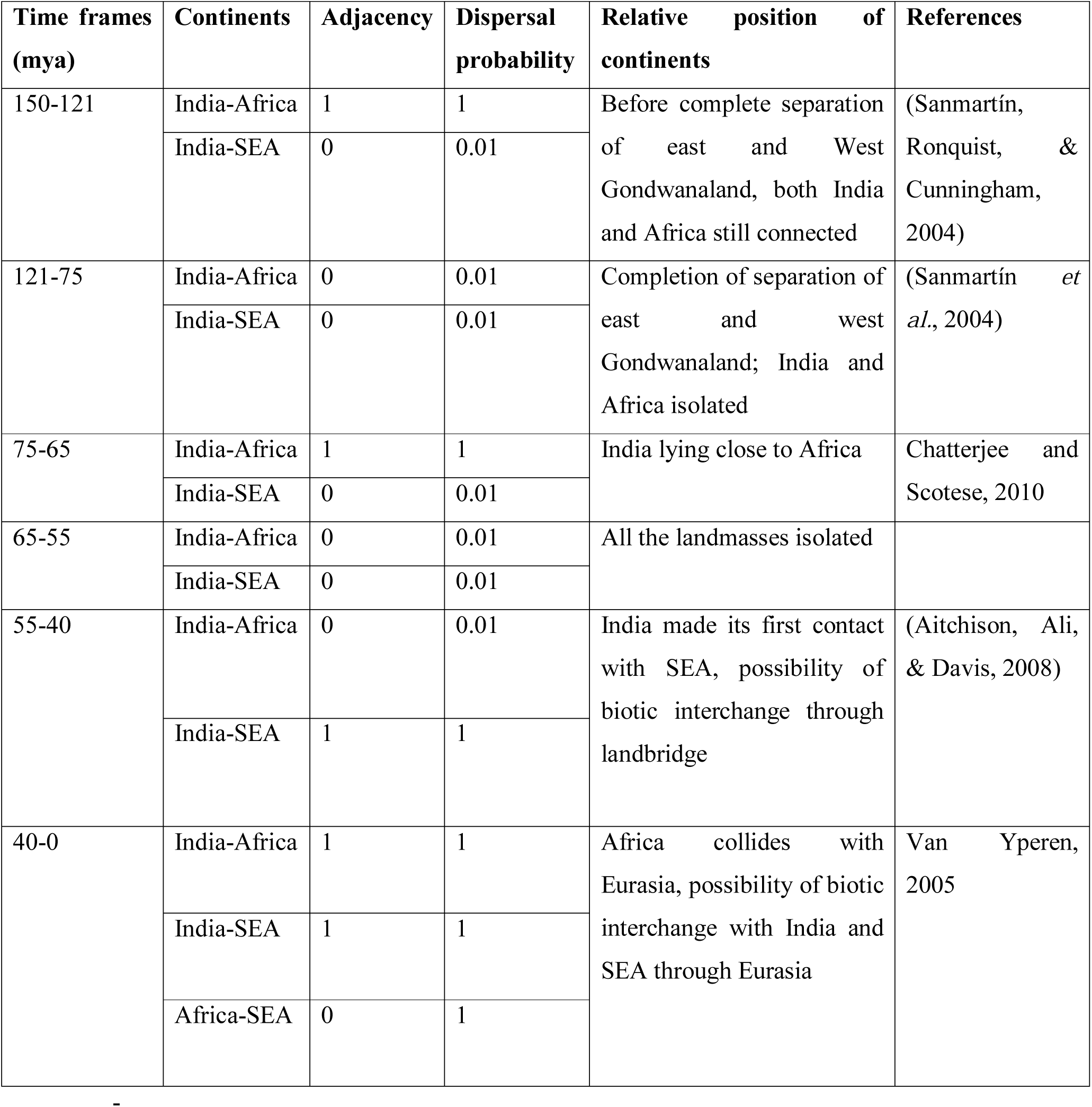
Adjacency and dispersal probability across different areas used in the ancestral range estimation analyses.

| Time frames (mya) | Continents | Adjacency | Dispersal probability | Relative position of continents | References |
| --- | --- | --- | --- | --- | --- |
| 150-121 | India-Africa | 1 | 1 | Before complete separation of east and West Gondwanaland, both India and Africa still connected | (Sanmartín, Ronquist, & Cunningham, 2004) |
|  | India-SEA | 0 | 0.01 |  |  |
| 121-75 | India-Africa | 0 | 0.01 | Completion of separation of east and west Gondwanaland; India and Africa isolated | (Sanmartín *et al.*, 2004) |
|  | India-SEA | 0 | 0.01 |  |  |
| 75-65 | India-Africa | 1 | 1 | India lying close to Africa | Chatterjee and Scotese, 2010 |
|  | India-SEA | 0 | 0.01 |  |  |
| 65-55 | India-Africa | 0 | 0.01 | All the landmasses isolated |  |
|  | India-SEA | 0 | 0.01 |  |  |
| 55-40 | India-Africa | 0 | 0.01 | India made its first contact with SEA, possibility of biotic interchange through landbridge | (Aitchison, Ali, & Davis, 2008) |
|  | India-SEA | 1 | 1 |  |  |
| 40-0 | India-Africa | 1 | 1 | Africa collides with Eurasia, possibility of biotic interchange with India and SEA through Eurasia | Van Yperen, 2005 |
|  | India-SEA | 1 | 1 |  |  |
|  | Africa-SEA | 0 | 1 |  |  |

**Figure A1.**
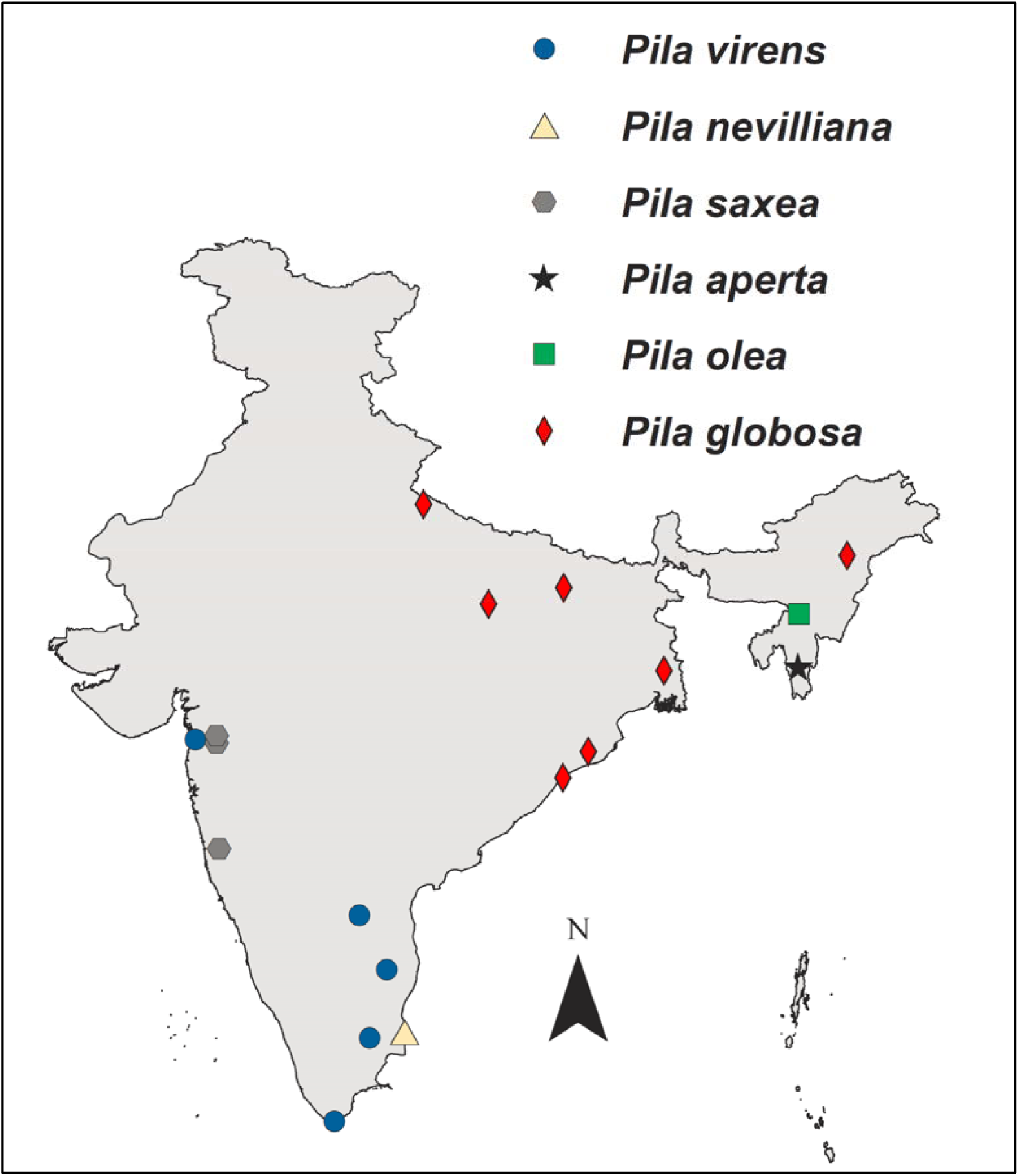
A map showing the location of individuals collected and amplified during the course of the study

**Figure A2.**
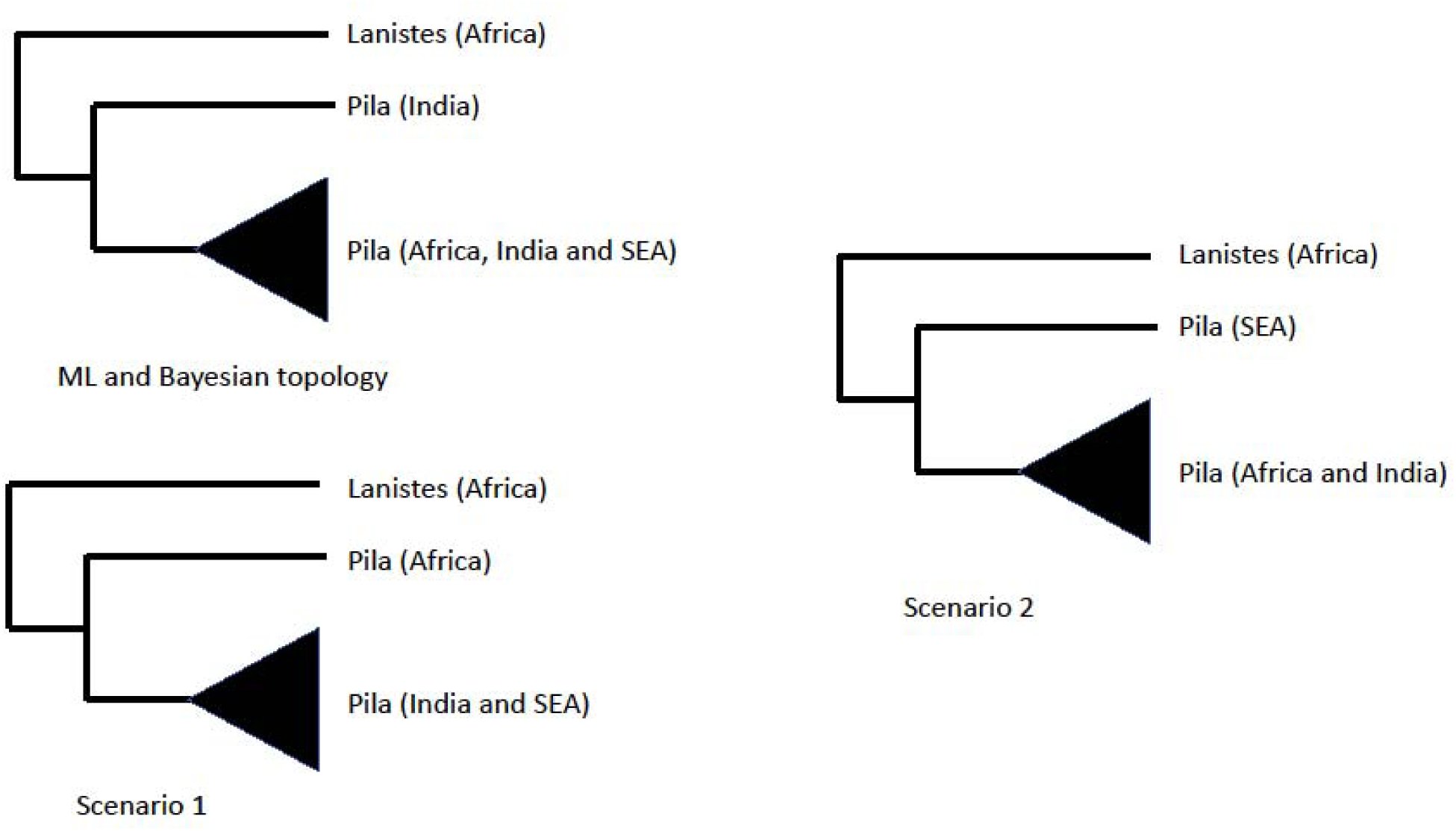
The different scenarios that were tested using AU test. The top left tree is the ML topology obtained from RAxML analysis and the tree summed over the posterior distribution of trees from the Bayesian analysis. In the bottom left scenario, African taxa, *Pila ovata* and *Pila speciosa* were constrained to be sister to the rest of the members of the *Pila* clade. In the scenario on the right, Southeast Asian taxa *Pila ampullacea, Pila conica, Pila polita* and *Pila aperta* were constrained to be sisters to the rest of the members of the *Pila* clade.

**Figure A3.**
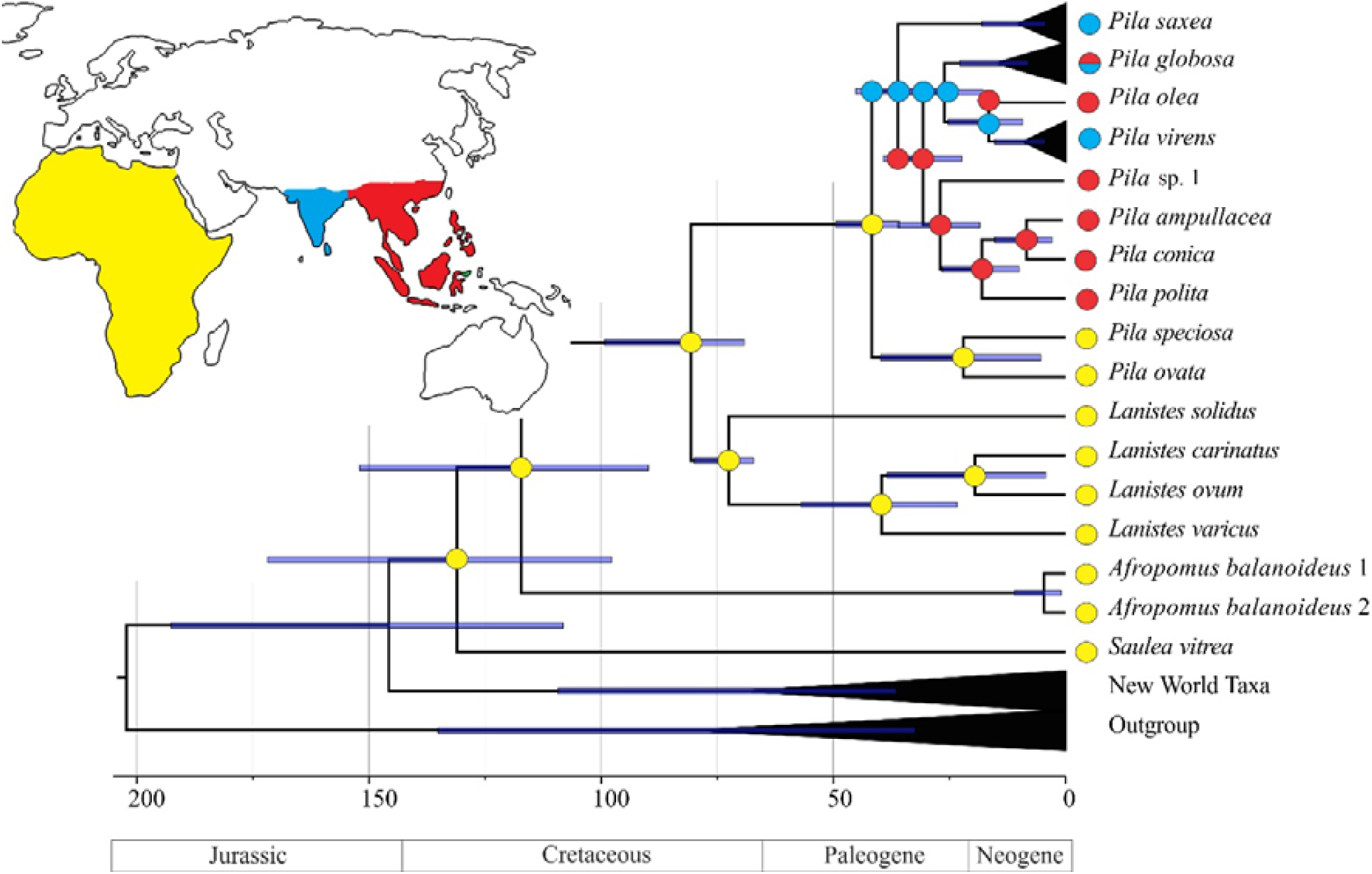
The MCC tree obtained from the BEAST analysis, where the African species of *Pila* were constrained to be sister to rest of the *Pila* species, with estimated ancestral ranges mapped on the nodes.

## Appendix B

In the current study two calibrations were used: a *Pila* fossil (Harzhauser *et al.*, 2016) excavated from Oman that dates back to Priabonian stage (37.8–33.9 mya) of Eocene epoch was used to calibrate the *Pila* MRCA. Arguably this is the oldest *Pila* fossil, since, older fossils than this could not be identified as *Pila* with certainty. Likewise, the *Lanistes* MRCA was calibrated with the oldest identified *Lanistes* fossil (Pickford, 2017) dating back to Maastrichtian (Late Cretaceous, 72.1–66 mya) deposits of Oman. We assumed that the MRCAs are slightly older than the fossils hence a gamma calibration was used (Ho & Phillips, 2009). In order to accommodate the uncertainty related to the age of the MRCAs, four independent analyses were carried out with different combinations of the shape (α) parameter, which alters the mode of the distribution that determines the age of the MRCA. The combinations were as follows: 1) *Pila* MRCA, α parameter=2, *Lanistes* MRCA α parameter =2; 2) *Pila* MRCA α parameter =2, *Lanistes* MRCA α parameter =4; 3) *Pila* MRCA α parameter =4, *Lanistes* MRCA α parameter =4; 4) *Pila* MRCA α parameter =4, *Lanistes* MRCA α parameter =). Similar strategy has been employed by other research groups previously in order to assign different levels of uncertainty to the calibrations and then select the best model (Klaus *et al.*, 2010). The scale (β) parameter was fixed at 2 for all the runs. Similarly, the offset was fixed at 37.8 mya and 66 mya for *Pila* MRCA and *Lanistes* MRCA respectively. Thereafter, Bayes factor test was employed in order to select the best model, which compares the marginal likelihoods of the runs. The marginal likelihoods were estimated using the harmonic mean estimator (Newton & Raftery, 1994; Suchard, Weiss, & Sinsheimer, 2001). The final run following the best model had the following calibration parameters: *Pila* MRCA, α parameter=4, β parameter=2, offset=37.8 mya; *Lanistes* MRCA, α parameter=4, β parameter=2, offset=66 mya.

